# Integration and harmonization of cell shape images for generative modeling

**DOI:** 10.64898/2026.02.07.704589

**Authors:** Alex Khang, Mark W. Young, Dilara Batan, Kristi S. Anseth

## Abstract

As cell imaging grows in scale, precision, and complexity, data integration and harmonization become increasingly important for studying cell–material interactions. Quantitative understanding of how cells respond to mechanical cues, such as substrate stiffness and topography, is often limited by differences in experimental conditions and imaging formats. This study presents a framework that combines compact, interpretable cell shape models with generative artificial intelligence to harmonize 2D and 3D immunofluorescent datasets within defined experimental contexts. By efficiently capturing morphology and associated biological features, the approach enables generation of realistic synthetic cells, including rare or intermediate phenotypes, to augment machine-learning analyses and support scalable in silico studies. This work advances data-driven investigation of cellular responses to biomaterial-derived mechanical cues.

## 1. INTRODUCTION

Currently, there is an extensive amount of cell imaging data that continues to expand on a daily basis as immunofluorescent imaging remains indispensable in illuminating protein abundance and structural organization. To achieve a collective and holistic understanding, efforts must be made toward integrating and harmonizing imaging data across the scientific community[1]. Here, we define data integration as an effort to establish a multidimensional dataset created from conceptually distinct data and harmonization as standardizing data gathered from different sources. Inspired by recent efforts[2,3], our approach is primarily focused on integrating and harmonizing data from two- and three-dimensional immunofluorescent images, and the quantitative data that can be derived from them. Specifically, we apply our approach to two representative cases: (1) single fibroblastic cells interacting with the surrounding matrix, and (2) dense epithelial cell monolayers characterized by abundant cell–cell interactions. Together, these cases capture the two major cellular organizations found in the body (mesenchymal and epithelial) and illustrate the versatility of our methodology across distinct biological contexts.

Imaging data can be directly used as feature vectors that describe single cells or a dense collection of cells. These feature vectors can be used to train artificial intelligence (AI) or machine learning (ML) models for various tasks including cell classification and generative modeling. However, this approach results in large data sizes and requires substantial computational power and time to train AI/ML models, often hours to weeks[4,5]. Dimensionality reduction techniques, such as principal component analysis (PCA), uniform manifold approximation and projection (UMAP), and autoencoders, can transform feature vectors into compact, yet meaningful, representations to accelerate model training[6]. Although PCA is generally interpretable, it often captures only linear relationships within the data and may not adequately preserve complex, non-linear patterns inherent in high-dimensional imaging data. In contrast, non-linear methods like UMAP and autoencoders are better at preserving intricate structures and clustering patterns but at the cost of reduced interpretability. The trade-offs between data compactness, information retention, and interpretability are key considerations when selecting dimensionality reduction strategies for downstream tasks in AI/ML applications involving cell imaging data.

Previously, quantitative shape models have been used as compact and interpretable representations of cell morphology derived from imaging data[7–9]. The use of cell shape modeling achieves dimensionality reduction naturally by representing cell boundaries using significantly fewer coefficients than the original boundary points. Recently, cell shape models have been used to integrate morphological and protein expression data derived from immunofluorescent images[10]. Building on this innovative work, we sought to develop efficient cell shape models for data integration, harmonization, and generative modeling of both 2D and 3D cells captured in immunofluorescent images (Fig. 1&2). Then, generative models, specifically β-variational autoencoders (β-VAEs), are trained to produce unique and biologically plausible synthetic cells based on features learned from the integrated and harmonized dataset (Fig. S.1&2). Importantly, generative modeling provides the ability to synthesize a diverse population of synthetic cells that reflect the underlying variability captured across many experiments, conditions, and research groups, rather than being restricted to selecting a limited number of discrete cells from a single dataset.

**Fig. 1.**
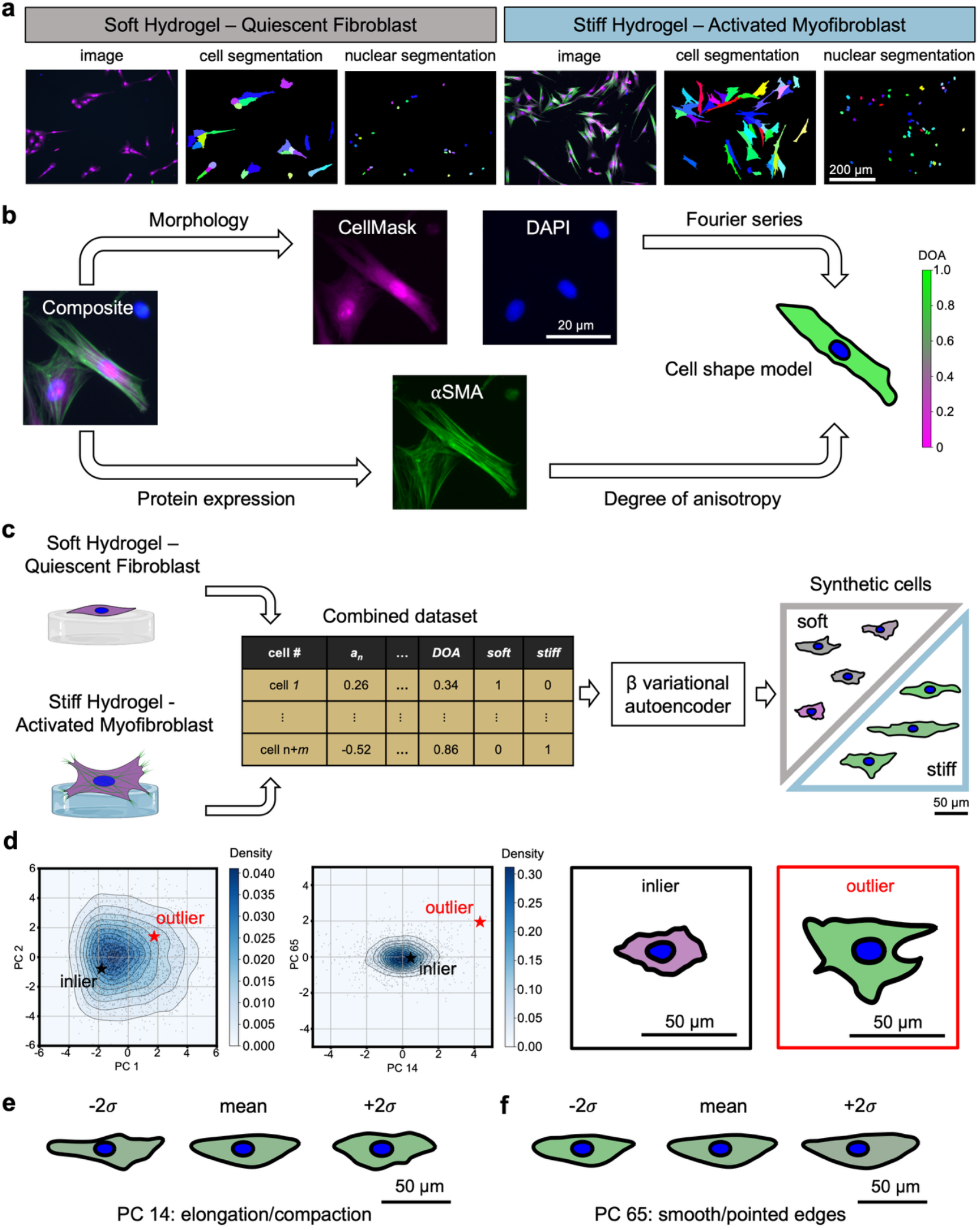
Data integration, harmonization, and generative modeling of 2D shape models derived from fluorescent images of valve interstitial cells (VICs). **a**, VICs seeded on soft hydrogels exhibit a quiescent fibroblast phenotype whereas VICs seeded on top of stiff hydrogels show an activated myofibroblast phenotype. Segmented nuclei are used as seed points for watershed segmentation of the cell cytoplasm. The scale bar applies to all images in this subpanel. **b**, Cell shape models integrate morphological and protein expression information and are achieved through Fourier series modeling of VIC body and nuclear segmentations colored by the corresponding degree of anisotropy (DOA) of intracellular alpha-smooth muscle actin stress fibers. The scale bar applies to all images in this subpanel. **c**, Data harmonization of different experimental groups is achieved through consistent Fourier series modeling and DOA computation for VICs seeded on soft and stiff hydrogels. One-hot encoding enables machine readable categorical features for every cell (e.g., being derived from the soft or stiff dataset). Data harmonization also enables the training of a single generative model, such as a β-VAE, that can synthesize unique but biologically plausible feature vectors of VICs seeded on soft or stiff hydrogels. **d**, Generative shape modeling facilitates the study of common (inlier, black star) and rare (outlier, red star) VICs as determined in principal component space. The outlier exhibits a low density value in the PC1 & PC2 plane (∼0.015) with the lowest density value occurring in the PC14 & PC65 plane (∼0). Cell shape models are produced from the generated feature vectors to visualize inlier and outlier VICs. The use of cell shape models and PCA allows for the visualization of eigenshapes along **e**, PC 14 and **f**, PC 65 axes to investigate properties indicative of rare VIC phenotypes. PCs 14 and 65 were physically interpreted as elongation/compaction and smooth/pointed edges, respectively. Stained images: green – *α*SMA, magenta – cell cytoplasm, blue – nuclei.

β-VAEs have been previously used to generate integrated instances of cells and their underlying organelles[4,5]. However, the use of β-VAEs to learn and generate efficiently parameterized cell shape models remains underexplored, despite their potential to accelerate *in silico* cell biology experiments for hypothesis generation and testing. To demonstrate the utility of generative shape modeling, we generate synthetic cells that lie on the outskirts of the training data distribution that may represent rare or underrepresented cell types, enabling their study in a controlled and scalable manner. Next, we utilize generative shape modeling to synthesize augmented training data for machine learning models to classify cell phenotypes based solely on morphological properties (Fig. S.3&4). Lastly, we showcase the usefulness of generative shape modeling in creating scientific illustrations that are on par with existing commercial and open-source selections (Fig. S.5&6).

## 2. MATERIALS AND METHODS

### 2.1 Valve interstitial cell isolation and culture on soft and stiff hydrogels

Valve interstitial cells (VICs) were isolated from porcine aortic valve leaflets obtained through dissection following previously published protocols[11]. Ten porcine hearts were sourced from Animal Biotech Industries, rinsed in Earle’s Balanced Salt Solution (Sigma) containing 1 µg/ml amphotericin B (Gibco), 50 µg/ml streptomycin (ThermoFisher), and 50 U/ml penicillin (ThermoFisher). Leaflets were enzymatically digested using 250 U/ml of type II collagenase (Worthington), dissolved in Earle’s solution, for 30 minutes to strip away the endothelial cell layer, which was subsequently discarded. The leaflet tissue was then incubated in a fresh collagenase solution for an additional 60 minutes under gentle agitation.

Following enzymatic digestion, the tissue suspension was vortexed for 5 minutes to promote VIC dissociation, and the resulting cell mixture was passed through a 100 µm cell strainer. The isolated VICs were then pelleted by centrifugation and resuspended in growth medium consisting of M199 (ThermoFisher) supplemented with 10% fetal bovine serum (FBS, ThermoFisher), 1 µg/ml amphotericin B, 50 µg/ml streptomycin, and 50 U/ml penicillin. Cells were expanded until they reached approximately 80% confluency, at which point they were detached using trypsin (ThermoFisher) and either passaged, frozen, or used in experimentation.

To ensure biological variability was minimized and to generate sufficient cell quantities for experimentation, VICs from all 10 porcine hearts were pooled. This approach helped to mitigate the effects of inter-animal variation and ensured consistency across experiments. Because individual valves yield a limited number of cells, pooling not only increased total cell yield but also enhanced reproducibility by averaging out variability from single animals. All hearts were processed using standardized protocols to reduce batch effects, and the n values reported refer to replicates derived from this pooled VIC population.

VICs were seeded on hydrogel substrates with tunable stiffness to model different mechanical environments: soft hydrogels (elastic modulus ∼2.8 kPa, n = 8) and stiff hydrogels (∼13.4 kPa, n = 4), as previously described (see Fig. 1a). Cells on soft matrices generally remained in a more quiescent state with reduced proliferation compared to those on stiffer, activation-promoting substrates. To ensure balance in training datasets, four additional soft hydrogel replicates were prepared to match cell numbers from the stiff condition.

Hydrogels were synthesized by co-polymerizing poly(ethylene glycol) photodegradable diacrylate (PEGdiPDA; MW ∼3,400 Da), poly(ethylene glycol) monoacrylate (PEGA; MW ∼400 Da), and acrylamide diethylene glycol – diethylene glycol – glycine – arginine – glycine – aspartic acid – serine – glycine (Ac-OOGRGDSG) in PBS using redox-initiated free radical polymerization[12]. Hydrogel formulations contained 7.0 wt% PEGdiPDA, 6.8 wt% PEGA, 5 mM Ac-OOGRGDSG, 0.2 M ammonium persulfate (APS), and 0.1 M tetramethylethylenediamine (TEMED). Hydrogels (∼100 µm thick) were cast onto acrylated glass coverslips and allowed to polymerize for 6 minutes before being washed and stored in PBS.

Stiff hydrogels were used as-is, while soft hydrogels were generated by exposing a separate set of stiff gels to 365 nm UV light at 10 mW/cm^2^ for 6 minutes, inducing photodegradation and reducing stiffness. VICs were seeded at a density of 15,000 cells/cm^2^ on 12 mm coverslips coated with either soft or stiff hydrogels. Cells were maintained for 72 hours in M199 medium supplemented with 1% FBS, penicillin-streptomycin, and amphotericin B before being fixed and stained for imaging.

### 2.2 Intestinal epithelial cell isolation and culture on photo-inducible hydrogels

Murine small intestinal crypts were isolated from Lgr5-eGFP-IRES-Cre_ERT2 mice. The isolated crypts were cultured as organoids by encapsulating them within growth factor reduced Matrigel (Corning) and culturing in stem media. Organoid stem media for routine culture was comprised of basal media supplemented with EGF (50 ng/mL, R&D systems), Noggin (100 ng/mL, Peprotech), R-spondin conditioned media (5% v/v/, Organoid and Tissue Modeling Shared Resource CU Anshutz), CHIR99021 (3 μM, Selleckchem), valproic acid (1 mM, Sigma), and N-acetyl cysteine (1 mM, Sigma). For normal passaging, organoids were recovered from Matrigel using ice-cold basal media and sheared (21-gauge needle) to remove dead cells. Organoid basal media was comprised of Advanced DMEM F12 (Invitrogen) that was supplemented with N2 and B27 supplements (1x each, Thermo Fisher Scientific), Glutamax (1x, Gibco), HEPES (10 mM, Gibco), and penicillin-streptomycin (1x). The resulting organoid fragments were encapsulated in Matrigel and cultured in stem media. Routine passaging of organoids was performed every 4 days as a splitting ratio of 1:4, and the medium was refreshed every 2 days. The University of Colorado Institutional Animal Care and Use Committee has approved the animal protocol (no. 00084) for this research.

Hydrogels for IEC culture were formed by a step-growth polymerization between an 8-arm poly(ethlyene glycol) (PEG) macromer end group functionalized with dibenzocyclooctyne (PEG-8DBCO) and an allyl sulfide bis(PEG-azide) crosslinker. To facilitate IEC interaction, adhesion, and growth on these hydrogels the peptides azide-GFOGER (1 mM) (sequence: azido lysine-GGYGG(GPP)_5_GFOGER(GPP)_5_), where O represents hydroproline, and azide-BM binder (0.5 mM) (sequence: azidobutyric acid ISAFLGIPFAEPPMGPRRFLPPEPKKP) were supplemented into the hydrogel formulation. Hydrogels were formed by pre-conjugating azide functionalized peptides with PEG-8DBCO in PBS for 10 minutes, after which crosslinker was added and the contents were vortex mixed. The hydrogels were formed as cylinders affixed to coverslips by casting the gels as droplets within a circular gasket (8mm diameter opening, 300 *μ*m height) placed on a sigmacoted glass slide after which a thiolated coverslip was overlaid onto the gasket which forced the hydrogel droplet to conform to the space within the gasket. Hydrogels were formed at 4 wt% total polymer and on stoichiometric equivalent ratios (DBCO:Azide). The gels were allowed to react for 30 minutes at room temperature, and reached a shear storage moduli of approximately 1.6 kPa.

For IEC culture as a monolayer atop these photoresponsive hydrogels, the surface of the gel was first passivated to spatially constrain the regions available for cellular growth. Briefly, the hydrogel surface was placed in contact with a solution containing PEG dithiol (3 wt%, 5 kDa, Jenkem) and the photoinitiator Lithium phenyl-2,4,6-trimethylbenzoylphosphinate (LAP) (3 mM) by inverting the hydrogel onto a gasket (1 mm thickness) containing a circular void (10 mm) containing 50 μL of the passivation solution. The hydrogel was allowed to equilibrate for 60 s in this solution before being lithographically irradiated (365 nm, 10 mW cm^-2^) using a collimated UV light (Thor labs) and a photomask (Frontrange Photomask) containing 800 μm chrome circles separated by 1 mm. In this process, the irradiated regions underwent a surface photoinitiated reaction, which we found prevented cell spreading.

IEC monolayers were formed by seeding a suspension of dissociated intestinal stem cells onto the surface of the hydrogel. Briefly, hydrogels (G’∼1.6 kPa, 1mM GFOGER, 0.5 mM BM binder) were allowed to equilibrate in basal media at 37°C for 30 minutes. Organoids were collected from Matrigel and were enzymatically dissociated by incubating in a solution of DNAse I (10 mg, StemCell Technologies), N-acetyl cysteine (1 mM, Sigma), and the rock inhibitor y-27632 (1 μM, SelleckChem) in TrypLE (1 mL, Thermo Fisher) for 8 minutes at 37°C, with mechanical agitation at 4 min and 8 min of incubation. The enzymatic dissociation was quenched by adding blocking media (1 mL FBS, 8 mL basal media), and the cell solution was filtered (40 μm cell strainer). The cell pellet was resuspended in growth media (at least 50 μL per gel to be seeded) at a cell density of 1E6 cells/mL. This cell suspension was seeded onto the hydrogel surface in a 50 μL droplet and allowed to adhere for 1 hr at 37°C, after which 0.6 mL of growth media was added.

Confluent monolayers were typically formed following 48-72 hr of growth after seeding dissociated IECs. At confluence, the monolayer bearing hydrogels were incubated in media containing 1 mM LAP and 10 mM PEG dithiol (5 kDa, Jenkem) in FluoroBrite media containing: N2, B27, Glutamax, HEPES, EGF, Noggin, R-spondin-CM, CHIR99021, and valproic acid for 1 hr at 37°C. Monolayer bearing hydrogels were lithographically irradiated through a photomask containing arrays of holes of 75 μm diameter separated by 50 μm using collimated UV light (Thorlabs, 365 nm, 5 mW cm^-2^, 90 s). Post irradiation, hydrogels were cultured in growth media. For IEC monolayer culture, growth media was comprised of basal media supplemented with EGF (50 ng/mL, R&D systems) WRN-conditioned media (50% v/v, Organoid and Tissue Modeling Shared Resource CU Anshutz), CHIR99021 (3 μM, Selleckchem), valproic acid (1 mM, Sigma), and N-acetyl cysteine (1 mM, Sigma). This media was supplemented with thiazovivin (2.5 μM, Selleckchem) during the first two days of monolayer culture.

### 2.3 Immunostaining and imaging

Samples were fixed in 4% paraformaldehyde (PFA; Electron Microscopy Sciences) for 20 min at room temperature, followed by two 5 min washes in PBS. Permeabilization was performed in 0.1% Triton X-100 (Fisher Scientific) in PBS for 20 min at room temperature. Non-specific binding sites were blocked for 1 h in 5% bovine serum albumin (BSA; Sigma-Aldrich) in PBS. Samples were incubated overnight at 4 °C with mouse anti–α-smooth muscle actin (αSMA) antibody (Abcam, Cat. no. ab7817; 1:200 in 5% BSA/PBS). The following day, primary antibody was removed and samples were washed twice in PBST (PBS + 0.5% Tween-20) and once in PBS. Secondary labeling was carried out for 1 h at room temperature in PBS containing goat anti-mouse Alexa Fluor 488 (ThermoFisher, Cat. no. A-11001; 1:200), DAPI (1:1000), and HCS CellMask Orange (Life Technologies; 1:5000). Samples were rinsed twice with PBST and once with PBS prior to imaging. Confocal imaging was performed on an Operetta high-content imaging system (PerkinElmer) using a 20x air objective. Images were acquired as 1360 × 1024 × 10 z-stacks with a lateral resolution of 0.49 µm/pixel and a 1 µm step size along the z-axis.

IEC monolayer samples containing swelling induced monolayer deformations were fixed at day 3 of growth using 4% PFA for 15 min at room temperature. The samples were permeabilized at room temperature for 15 minutes using 0.1% Triton X100, blocked at room temperature for 1 hr using a solution of 5% BSA. Samples were incubated overnight at 4 °C in a solution of 5% BSA containing DAPI (1:1000) and AlexaFluor555 conjugated phalloidin (1:300, Cell Signaling Tech). Samples were washed in PBS (3x10 min) and imaged. Imaging was performed using a Nikon AXR laser scanning confocal microscope operating in resonance scanning mode with a 20x water immersion objective. Images were acquired with a 0.31 µm/pixel lateral resolution and a 0.31 µm step size along the z-axis.

### 2.4 Segmentation, centering, and alignment

The cell boundary of VICs were segmented using a marker-based watershed algorithm, implemented in Harmony High-Content Imaging and Analysis Software (PerkinElmer). Nuclear segmentations served as markers to delineate individual cells. Although this approach was carried out using Harmony, similar marker-based watershed segmentation techniques are widely available in platforms such as MATLAB, Python, ImageJ, and CellProfiler. Segmentations were manually reviewed for quality. Cells exhibiting segmentation artifacts—such as incomplete boundaries or inclusion of adjacent cells— were excluded from further analysis. Additionally, cells undergoing division, as evidenced by split nuclei, were removed. Only cells entirely contained within the field of view (i.e., not intersecting the image boundaries) were retained. For morphological quantification, the pixel coordinates defining segmented cell and nuclear boundaries were converted to Cartesian coordinates by multiplying pixel positions by the image resolution (0.49 µm/pixel). To normalize for position and orientation, each cell and nucleus was translated and rotated such that their centroids were aligned at the origin and their principal axes were oriented along the x-axis. The principal axis was defined as the direction capturing the greatest variance in boundary point distribution, determined via principal component analysis (PCA) of the Cartesian coordinates. In addition, the orientation was adjusted (e.g., flipping horizontally and/or vertically where necessary) so that most of the cell or nuclear area lay in the first quadrant. These shape-preserving transformations standardized the spatial distribution of major features, such as protrusions, across all cells to facilitate consistent morphological comparisons.

Fluorescent images of IEC monolayers contained channels corresponding to cell membrane (Factin) and cell nuclei (DAPI). These stains were custom processed to facilitate the 3D segmentation of IECs using the FIJI plugin LiMe Seg. Briefly, using ImageJ FIJI, the nuclei channel was blurred using a 3D gaussian blur filter (sigma = 3) and binarized (otsu). Next, the Factin channel was processed using a 3D gaussian blur filter (sigma = 3), followed by a rolling ball background subtraction (Radius = 10 pixels). The resulting image was binarized (Li). To generate seed points for use in LiMe Seg, the binarized Factin channel was subtracted from the binary nuclei channel, and the resulting image was used to calculate a 3D normalized Euclidian distance map (EDM) (FIJI). The EDM image was blurred using a 3D gaussian blur filter (sigma = 4) and then centroid positions of the nuclei center were determined using a 3D maxima finder (FIJI). These coordinates were used to draw circular ROIs (Radius = 5 pixels) that served as seed points for LiMe Seg. LiMe Seg (FIJI) was used to segment individual IECs using the Factin channel and the generated seed points. Resulting point clouds were exported from FIJI and used to generate IEC surface meshes.

The raw IEC surface meshes were centered on their centroid (Fig. S.7A&B). To ensure that IEC pose did not influence morphological analysis, IECs were rotated such that their basal surface was parallel with the x-y plane (Fig. S.7B&C). This was accomplished by first identifying the IEC’s basal surface which was denoted by facets whose surface normal points downward as defined by having a *z*-value less than -0.8 (red arrows in Fig. S.7B which are not drawn to scale). Then, an average basal surface normal was computed by averaging all the surface normal of the facets that comprise the basal surface (blue arrow in Fig. S.7B which is not drawn to size). Next, the IEC surface mesh is rotated such that the average basal surface normal is aligned along the global negative z-direction defined by the vector [0, 0, -1] (Fig. S.7C). Lastly, the IEC is rotated along the x-y plane so that the largest volume of the shape is in quadrants with positive x-values to ensure consistent morphological analysis across shapes and to reduce effects arising from pose and orientation (Fig. S.7D). Non-deformed and deformed IECs were determined based on the height or z-component of their centroid with centroids lower than z = 10 um being labeled as non-deformed and those above being labeled as deformed (Fig. 2E).

**Fig. 2.**
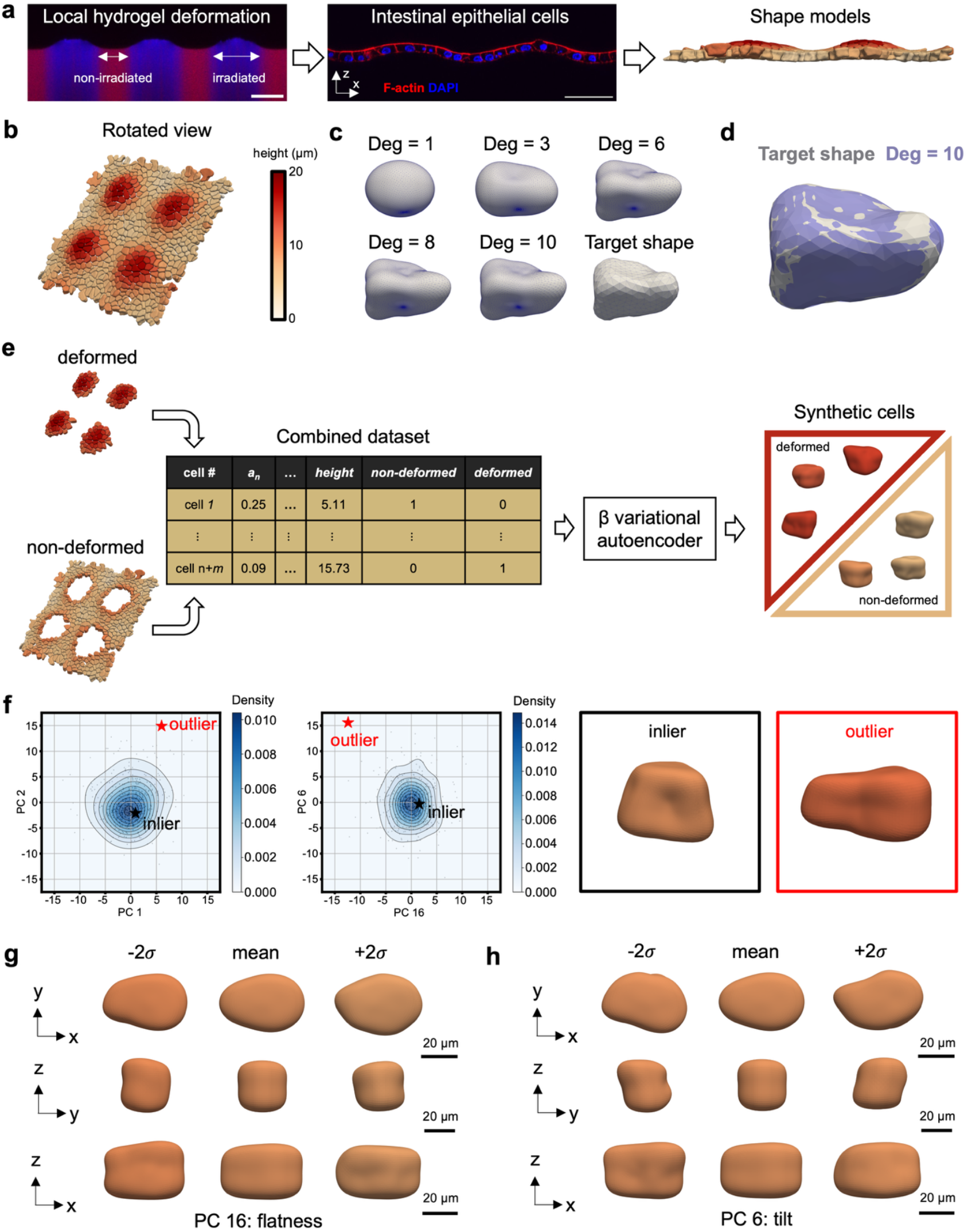
Data integration, harmonization, and generative modeling of 3D shape models derived from fluorescent images of intestinal epithelial cells (IECs). **a**, Ally sulfide crosslinked photoresponsive poly(ethylene glycol) hydrogels are equilibrated in medium supplemented with photoinitiator and thiol functionalized molecules and lithographically irradiated using collimated UV light to induce degradation (blue) induced swelling onto the top surface (left panel). This process is cytocompatible and can be used to pattern local deformations within confluent monolayers formed from IECs grown on the gel surface prior to inducing swelling (middle panel). Individual IECs are segmented to achieve 3D surface meshes (right panel). **b**, Three-dimensional visualization of the IEC monolayer surface meshes colored by height. Height is determined by the z-component of the cell centroid. **c**, Spherical harmonics (SPHARM) are used to model IECs and converge to the target shape with increasing SPHARM degrees. **d**, A degree of 10 was determined to be sufficient to model IECs as evidenced by the quality of fit between the target shape and SPHARM model. **e**, Data harmonization of deformed and non-deformed cells are achieved through consistent SPHARM modeling and one-hot encoding of IECs. One-hot encoding enables machine readable categorical features for every cell (e.g., deformed vs non-deformed). Data harmonization also enables the training of a single generative model, such as a β-VAE, that can synthesize unique but biologically plausible feature vectors of deformed and non-deformed cells. **f**, Generative shape modeling facilitates the study of common (inlier, black star) and rare (outlier, red star) IECs as determined in principal component space. The outlier exhibits a low density value in the PC1 & PC2 plane (< 0.002) with the lowest density value occurring in the PC16 & PC6 plane (∼ 0). Cell shape models are produced from the generated feature vectors to visualize inlier and outlier IECs. The use of cell shape models and PCA allows for the visualization of eigenshapes along **g**, PC16 and **h**, PC6 axes to investigate properties indicative of rare IEC phenotypes. PCs 16 and 6 were physically interpreted as flatness and tilt, respectively.

### 2.5 Modeling 2D segmentations using Fourier series

To quantitatively characterize VIC body and nuclear morphology, we applied a Fourier-based shape analysis method that provides compact mathematical descriptions of boundary geometry[13]. Cell and nuclear contours were parameterized using two functions, *x(θ)* and *y(θ)*, which map each normalized arc-length position θ ∈ [0, 2π] along the shape boundary to corresponding Cartesian coordinates. This normalization ensures a one-to-one correspondence between each point on the boundary and a unique angle θ, effectively reparameterizing the original boundary onto a unit circle. Each function was expressed as a truncated Fourier series with n harmonics:

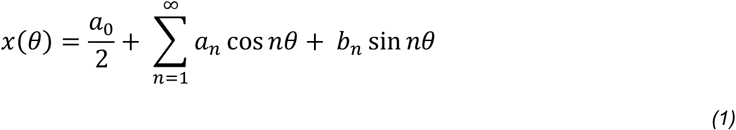

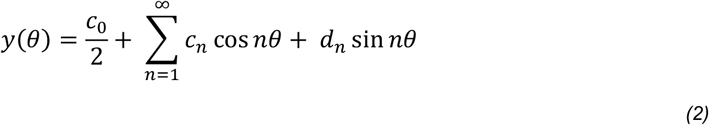

The Fourier coefficients (*a*_*0*_, *a*_*n*_, *b*_*n*_, *c*_*0*_, *c*_*n*_, and *d*_*n*_) were computed by solving an overdetermined system of linear equations constructed from uniformly sampled boundary points. These systems were solved in matrix form using least squares regression to obtain the best-fit coefficients for a given number of harmonics. As the number of harmonics increases, the reconstructed boundary more closely approximates the original shape, enabling high-fidelity shape representation (Fig. 1d). For this study, n = 15 and 5 for the cell body and nuclei respectively as determined by the observation at which increasing the number of Fourier harmonics no longer resulted in noticeable decreases in residuals sum squares (RSS), Bayesian Information Criterion (BIC) or Akaike Information Criterion (AIC)[10]. In total, VIC shapes were modeled using 84 unique features (62 coefficients for the cell body and 22 coefficients for the nuclei). The resulting set of Fourier coefficients encapsulates both size and geometric features of each cell and nucleus, serving as a compact, data-driven representation for downstream morphological analysis.

### 2.6 Modeling 3D segmentations using spherical harmonics

We employed spherical harmonics to model IEC shapes[13]. First, IEC surface meshes were subjected to spherical parameterization to create a continuous, uniform mapping from the original surface to the surface of a unit sphere[7]. This was accomplished by interpolating the original shape over a spherical grid with a resolution of 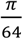 along the polar (0 ≤ *θ* ≤ *π*) and azimuthal (0 ≤ *φ* < 2*π*) angles using nearest neighbors. With the inclusion of north and south poles, this resulted in a total of 8,194 vertices in the interpolated mesh. Ultimately this achieves a one-to-one mapping between Cartesian and spherical coordinates for each vertex on the surface mesh (*x*(*θ, φ*), *y*(*θ, φ*), *z*(*θ, φ*)). Then, the interpolated surface mesh was fit to a set of SPHARM basis functions 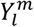 with the following form:

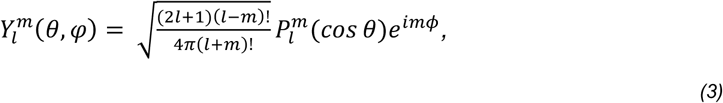

where *l* and *m* denote the degree and order of the SPHARM, respectively, and 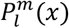 is the Legendre polynomial as follows:

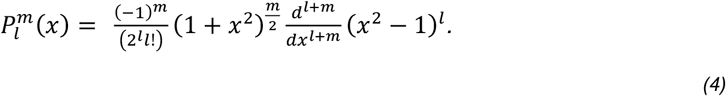

Each Cartesian *x, y, z* coordinate was independently represented with SPHARM as such:

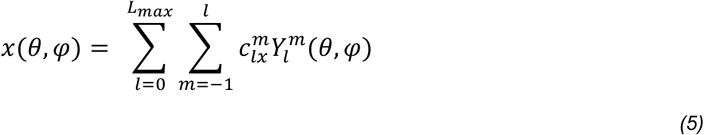

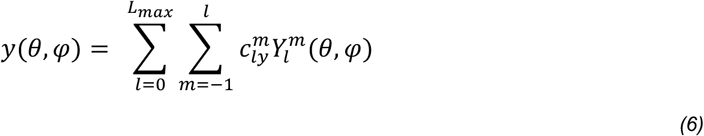

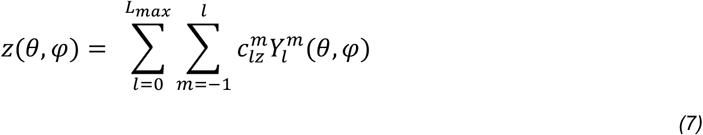

where 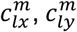, and 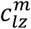 are the SPHARM coefficients for the *x, y, z* coordinate components, respectively. The SPHARM coefficients were determined at a user specified maximum degree *L*_*max*_. For this study, *L*_*max*_ = 10 was sufficient to reconstruct IEC surface meshes based on the observation that increasing the SPHARM degree past 10 no longer resulted in appreciable decreases to the RSS, BIC, or AIC (Fig. S.8). This resulted in 121 coefficients for 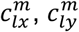, and 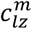 which contained real and imaginary components. In total, IEC shapes were modeled using 726 unique features (121 coefficients x 3 coordinate components x 2 real/imaginary components).

### 2.7 Computing degree of anisotropy of *α*SMA stress fibers in VICs

To quantitatively assess the degree of anisotropy (DOA)[10] and organization of *α*SMA stress fibers, we first computed coherence, a scalar descriptor of local image anisotropy. This method provides an objective measure of stress fiber discreteness and orientation, based on previously established gradient structure tensor analysis. First, pixel-wise image gradients in the *x-* and *y*-directions (*I*_*x*_ and *I*_*y*_, respectively) were calculated using the Sobel-Feldman operator, which applies convolution with the following 3×3 kernels:

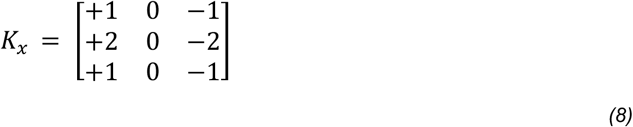

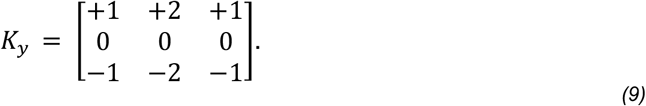

Image gradients were computed via two-dimensional convolution:

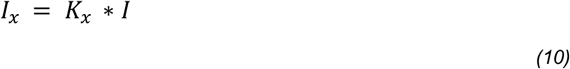

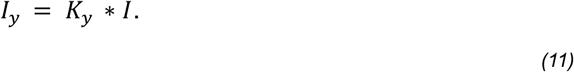

where *I* is the raw fluorescence image and * denotes convolution. The gradients were used to construct a structure tensor *S* for each pixel, which captures local intensity variations and orientation:

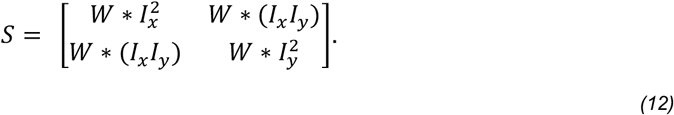

To suppress noise and incorporate neighboring information, each component of the structure tensor was smoothed with a Gaussian kernel *W* with standard deviation of 4 pixels, corresponding to the approximate diameter of *α*SMA stress fibers in our images. The standard deviation can be tuned depending on image resolution and the size of the structure of interest. Eigenvalue decomposition of the structure tensor yielded two eigenvalues (*λ*_1_, *λ*_2_) and associated eigenvectors (***e***_1_, ***e***_2_) which describe the magnitude and direction of local intensity changes. A high degree of anisotropy is indicated by *λ*_1_>>*λ*_2_, whereas *λ*_1_=*λ*_2_ suggests isotropic regions or homogeneous intensity. In the limiting case where *λ*_2_ → 0 and *λ*_1_ > 0, the image is locally one-dimensional, corresponding to perfectly aligned structures. To quantify anisotropy at each pixel, we computed coherence (*c*) as:

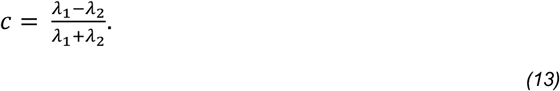

Coherence ranges from 0 (no preferred direction) to 1 (perfect alignment). A single scalar descriptor of cellular *α*SMA organization was obtained by computing the median coherence value across all pixels within each segmented cell body, yielding DOA (Fig. 1b). Traditionally, *α*SMA stress fiber presence and organization were assessed qualitatively to classify fibroblast activation states. In contrast, DOA provides a robust, quantitative alternative.

### 2.8 Generative shape modeling using β-variational autoencoders

Raw data were imported from an Excel spreadsheet containing Fourier shape descriptors of VICs or SPHARM shape descriptors of IECs. Each row represented a single cell with columns divided into shape model coefficients, DOA or height, and two one-hot encoded categorical features located in the final two columns that represent soft/stiff culture conditions for the VICs or non-deformed/deformed for the IECs. The numeric portion of the data was normalized feature-wise by subtracting the mean and dividing by the standard deviation. The normalized numeric data were then concatenated with the one-hot encoded categorical variables to form a composite input matrix, which was subsequently converted to a PyTorch tensor for model training.

We implemented a fully connected β-variational autoencoder (β-VAE) using PyTorch (Extended Data Fig.1). The encoder consisted of two hidden layers: a linear transformation from input dimension *n* to 8*n* units followed by a ReLU activation and a second transformation to 4*n* units followed by another ReLU activation. Two parallel linear layers then projected the intermediate representation into mean (*μ*) and log-variance (*logσ*^2^) vectors of a 15-dimensional latent space. Sampling was performed via the reparameterization trick. The decoder mirrored the encoder with two hidden layers and ReLU activations, followed by a linear output layer that reconstructed the original input dimension *n* (Extended Data Fig.2). The latent dimensionality and hidden layer sizes were empirically chosen to balance model expressiveness with reconstruction fidelity. The β-VAE was trained for 500 epochs using the Adam optimizer (learning rate = 1e-3) on the full dataset. The loss function was a weighted sum of mean squared error (MSE) reconstruction loss and the Kullback–Leibler divergence (KLD) between the approximate posterior and a unit Gaussian prior. To limit latent space regularization during training and preserve fine-grained structure in the learned distribution, we employed a small β of 1e-6 on the KLD term and 1-β on the MSE term. Model weights were initialized with PyTorch’s default settings, and all random seeds were fixed to ensure reproducibility.

After training, the decoder was used to generate synthetic cells by sampling latent vectors *z*∼𝒩(0, ***I***). The output of the decoder was split into normalized numeric and categorical parts. The numeric features were denormalized using the original training set statistics (mean and standard deviation). For the VIC dataset, a post-processing step clipped the generated DOA values to the range [0, 1] to enforce biologically plausible bounds. For the two categorical variables, one-hot vectors were reconstructed by applying an argmax operation and converting the result back into one-hot encoding.

To assess the fidelity of the synthetic data, we compared its statistical structure to that of the real dataset using Spearman correlation heat maps (Fig. S.9). We computed pairwise Spearman correlations among all features for both the real and generated datasets and correlation matrices were visualized as heatmaps. We compare heatmaps from the training dataset, β-VAE synthetically generated dataset, and a synthetic dataset produced via a naïve approach. In the naïve approach, a kernel density estimate is computed for each component of the cell feature vectors. Random values are then drawn from each kernel density estimate to achieve a synthetic cell feature vector. Furthermore, to verify that the β-VAEs were generating novel cell shapes rather than simply reproducing examples from the training data, we visualized three representative synthetically generated VICs and IECs alongside their closest real counterparts from the training dataset (Fig. S.10).

### 2.9 Identification of outliers in synthetically generated data

To identify statistical outliers among synthetic and real cell shape representations, we employed a multivariate outlier detection approach based on principal component analysis (PCA) followed by Mahalanobis distance computation in the reduced feature space. First, the one-hot encodings were removed from the training data and the remaining feature vectors were standardized by subtracting the mean and dividing by the standard deviation of each feature. PCA was then performed on the covariance matrix of the standardized training data using eigen decomposition. Principal components (PCs) were sorted by explained variance, and the minimum number of PCs required to capture ≥95% of the total variance was retained (*K* = 66 for the VIC dataset and *K* = 86 for the IEC dataset). This transformation reduced the dimensionality while preserving relevant information. Both the training data and the generated synthetic data were projected into this reduced PC space using the same eigenvectors. Within the reduced PC space, the Mahalanobis distance of each synthetically generated cell was computed relative to the training data distribution. This distance measured how many standard deviations a point lies from the multivariate mean, accounting for feature correlations. The Mahalanobis distance *D*_*M*_ is given by:

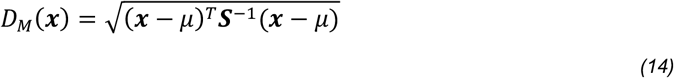

where ***x*** is a generated feature vector in PC space, *μ* is the mean of the training dataset in PC space, and ***S***^−1^ is the inverse of the training set covariance matrix in PC space. A synthetically generated outlier was defined using the 95^th^ percentile of the Chi-square distribution with *K* degrees of freedom. That is, any generated cell feature vector ***x*** with 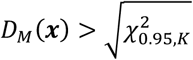 was classified as a statistical outlier.

### 2.10 Generation of eigenshapes to investigate properties of rare cells

For each identified outlier in the generated dataset, we quantified the contribution of individual principal components to its Mahalanobis distance. The squared contribution of each PC was computed by:

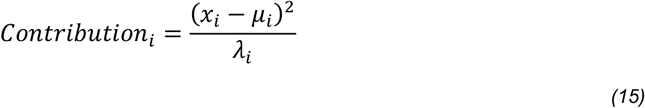

where *x*_*i*_ is the coordinate of the outlier in PC space, *μ*_*i*_ is the mean of the training data also in PC space, and *λ*_*i*_ is the corresponding eigenvalue. Here, *i* denotes which PC these values are being computed for. The PCs contributing most to the outlier status were ranked by their normalized contributions and visualized in 2D KDE plots (Fig. 1D&2F). To support interpretability, the training data was visualized along PCs 1 and 2 and in the top two contributing PCs (PC 14 & 65 for the VIC dataset, PC 16 & 6 for the IEC dataset). Kernel density estimation (KDE) was applied using Gaussian kernels to estimate the underlying distribution in 2D, and outliers were highlighted with star markers. These steps enabled both quantitative detection and qualitative visualization of outliers, revealing which modes of variation (principal components) contributed most significantly to anomalous data points. To investigate the morphological characteristics that contribute towards synthetically generated outliers, we computed eigenshapes along the top two contributing PCs by taking the average of the PC scores of the training dataset, systematically altering the average value of the PC axis of interest from -2*σ* to 2*σ*, transforming the PC scores back to original feature vectors, and then visualizing shapes using Fourier series or SPHARM expansion (Fig. 1E,F & 2G,H).

### 2.11 Generative shape modeling to augment training of machine learning models

To investigate the utility of using generative cell shape modeling to augment training of machine learning models, we trained several multivariable logistic regression models on a combination of real and synthetic data to predict whether VICs were from the soft or stiff hydrogel culturing condition based on their Fourier series coefficients (Fig. S.3A). Likewise, this process was repeated for IECs in which their SPHARM coefficients were used to predict whether they were non-deformed or deformed (Fig. S.4A). It is important to note that centering and alignment of the IECs ensures that the models learn only morphological features and do not make their predictions based on position (i.e., height) nor pose. This allows us to gauge the extent at which morphology alone is predictive of IEC class. We trained models using 100% real data, 50% real data, and 50% real data + varying levels of β-VAE generated synthetic data derived from the 50% real data to assess whether data augmentation could increase model accuracy compared to training using only real data. To visually and quantitatively assess the performance of the logistic regression models, we plotted receiver operating characteristic (ROC) curves (Fig. S.3B&4B) and computed the corresponding area under the curve (AUC, Fig. S.3C&4C).

### 2.12 Generative shape modeling for scientific illustrations

We demonstrate the proof of concept for using generative shape modeling to produce scientific illustrations of 2D fibroblast shapes (Fig. S.5A). We generated synthetic VICs using our trained β-VAE and compare the shapes visually to fibroblast illustrations gathered from open-source (NIH BioArt) and commercial (Biorender) sources. Additionally, to demonstrate proof of concept of using shape models for data harmonization, we subjected the NIH BioArt and Biorender shapes to the same Fourier series modeling performed on our VIC dataset and trained an additional β-VAE to generate novel shapes derived from the three separate sources (Fig. S.5B). Moreover, to showcase the ability to customize β-VAE cell illustrations, we generated several examples in which the maximum limit of Fourier series is tuned to achieve varying levels of coarseness and fineness in the illustrations. Lastly, we demonstrated the ability to tune the color for the stroke and fill of the cell body and nuclei, which further increases the customizable nature of these illustrations (Fig. S.6).

## 3. RESULTS

### 3.1 Generative modeling of VICs seeded on top of soft and stiff hydrogel surfaces

For our first test case, we demonstrate the utility of cell shape models using 2D segmentations of valvular interstitial cells (VICs) derived from the aortic valve using publicly available immunofluorescent images[14] (Fig. 1A). Porcine VICs were seeded on soft (∼2.8 kPa) and stiff (∼13.4 kPa) hydrogel surfaces, which promoted a quiescent fibroblast and activated myofibroblast phenotype, respectively. Activated myofibroblasts play a critical role in wound healing and show a characteristic increase in expression of alpha smooth muscle actin (*α*SMA) stress fibers, are highly contractile, and exhibit elevated extracellular matrix secretion and remodeling[15]. Cell body and nuclear segmentations were used to develop cell shape models using a Fourier series[13]. The degree of anisotropy (DOA)[10] of the *α*SMA stress fibers for each cell was computed and integrated into the cell shape model to enable the representation of morphology and protein expression simultaneously (Fig. 1B). Next, we demonstrated that consistent representation of VICs from different experimental conditions (as well as from different sources (Fig. S.5)) using shape modeling enabled data harmonization and subsequent training of a single β-VAE that could synthesize unique but biologically plausible VICs (Fig. 1C). We identified and visualized a representative inlier and outlier produced by the β-VAE, based on the training data projected into principal component space (Fig. 1D). We further generated eigenshapes corresponding to the two principal components that had the greatest influence on distinguishing the representative outlier from inliers (Fig. 1E). This analysis revealed that the outliers contained a unique combination of morphological characteristics, such as compactness and smooth and pointed edges, which are not typically observed in VICs with high DOA values derived from culturing on stiff hydrogel substrates[10]. Interestingly, the outliers had characteristics of both quiescent fibroblasts (compactness, smooth edges) and activated myofibroblast (high DOA value, pointed edges indicative of protrusion formation) that are reminiscent of a proto-myofibroblast[15].

### 3.2 Generative modeling of IECs seeded on top of hydrogel surfaces with photoinducible topology

Next, we extend the use of cell shape modeling to represent 3D segmentations of dense cell monolayers to understand the dependency of their morphology on the underlying substrate topology (Fig. 2A). This dataset was generated by culturing intestinal epithelial cells (IECs) as a monolayer on the surface of a photo-responsive allyl sulfide crosslinked hydrogel that allowed for user-controlled changes in the topographical features in the epithelium (e.g., undulations similar to in vivo morphology). Changes in the IEC monolayer were dictated by controlled lithographic irradiation during culture to induce deformations and dynamically alter the epithelial curvature. These changes were captured by immunofluorescent images of f-actin and segmented to create 3D surface meshes of individual IECs within the confluent monolayer.

These 3D cell shape models enabled the integration of additional information such as cell height in the monolayer (Fig. 2B). Individual IECs were modeled using spherical harmonics (SPHARM) which converge to the target shape when the degree is sufficient (Fig. 2C). We found that a SPHARM degree of 10 was sufficient to model IEC shapes (Fig. 2D). Similarly to the VIC dataset, consistent shape modeling enabled the harmonization of cells localized within deformed and non-deformed regions of the hydrogel to train a β-VAE capable of generating synthetic but biologically plausible IECs (Fig. 2E). We note that data harmonization is not limited to this case and may be helpful to standardize and combine datasets from different experimental conditions and research groups. The β-VAE generated inliers and outliers (Fig. 2F). The representative outlier’s centroid was positioned approximately at 10 *μ*m in height and appeared to lie midway between deformed and non-deformed cell morphologies. The generated eigenshapes associated with the two principal components most influential in distinguishing the representative outlier from inliers revealed that a distinctive combination of flatness and tilt contributed to its outlier status (Fig. 2G&H).

### 3.3 Using generative modeling to augment training data for machine learning

Another valuable application of generative shape modeling is data augmentation for training machine learning models (Fig. S.3&4), which can be beneficial in data-scarce scenarios. To demonstrate the utility of this approach, we generated synthetic VICs and IECs and evaluated how supplementing the training dataset with these synthetic cells affected model accuracy compared to training with real data alone. For this task, we employed multivariable logistic regression models to predict whether VICs were from the soft or stiff culture condition based solely on their Fourier coefficients, which implicitly contains their morphological properties (Fig. S.3A). Likewise for the IEC dataset, we trained multivariable logistic regression models to predict whether an IEC was a deformed or non-deformed cell based solely on their SPHARM coefficients (Fig. S.4A). We report that for both cases, data augmentation can increase model accuracy, although accuracy did not necessarily scale with the amount of synthetic data used for augmentation (Fig. S.3B-C&4B-C).

### 3.4 Applications in scientific illustration

Lastly, we used generative shape modeling to create unique and diverse cellular shapes for scientific illustration (Fig. S.5&6). Our representations were on par with open-source (NIH BioArt) and commercial selections (Biorender), and we further demonstrated the usefulness of our approach in data harmonization by parameterizing the NIH BioArt and Biorender images using the same cell shape modeling framework used for the VIC dataset. For a further demonstration, we trained a β-VAE to learn features from all three sources to produce novel cell illustrations (Fig. S.5) that were highly tunable in terms of color as well as coarseness/fineness of cellular details (Fig. S.6).

## 4. DISCUSSION

Herein, we present an approach to efficiently integrate and harmonize cellular data derived from immunofluorescent images. One key advantage of using efficient cell shape models is the significantly reduced training time for generative AI, requiring minutes rather than the hours or even weeks when training on full-resolution images[4,5]. When combined with generative modeling, the approach can accelerate the investigation of understudied or rare cell types (e.g., proto-myofibroblasts[15], Merkel cell carcinoma[16], etc.). It is important to note that our approach is highly flexible and can facilitate different cell shape modeling methods such as VAMPIRE[17] or Zernike polynomials[18]. Furthermore, we acknowledge that alternative model architectures such as transformers, GANS, and autoregressive models can also be used for generating synthetic cells.

Our investigation supports that cell shape models are a practical tool for efficient data integration, harmonization, and generative modeling. Looking forward, these capabilities are a step toward integrating diverse datasets (e.g., morphology + spatial omics[19]) harmonized from diverse sources (Fig. S.5) with utility in investigating spatiotemporal changes in cell state within developing or maturing organoids and tissues. Importantly, these types of analyses will help researchers elucidate relationships between cell morphology and state/function to potentially infer cellular behaviors based solely on shape. Altogether, these advances should open the door to hypothesis generation and experimentation using virtual cells[20,21], enabled by generative modeling to investigate single and collective cell responses. Going forward, generative cell shape modeling should be combined with cell-signaling, agent-based, and continuum models to simulate cellular function across several length scales.

## 5. CONCLUSIONS

We demonstrated that generative modeling of cell shape provides a powerful framework for representing, synthesizing, and analyzing cellular morphology in response to distinct material environments. Using 2D and 3D segmentations of VICs and IECs cultured on soft, stiff, and photoinducible hydrogel surfaces, we showed that harmonized shape models can capture key phenotypic adaptations to substrate stiffness and topography while enabling the generation of biologically plausible synthetic cells via β-VAEs. These models revealed unique morphological outliers, including transitional proto-myofibroblasts and intermediate IEC shapes, highlighting how material cues influence cellular morphology.

Furthermore, synthetic cells generated from these models can augment training datasets for machine learning and produce versatile scientific illustrations, demonstrating the practical utility of generative approaches in data-scarce or multi-source scenarios. By integrating cellular shape with material context, our results highlight the potential for predictive modeling of cell–material interactions, enabling researchers to investigate how substrate mechanics and topology influence cell behavior. Generative cell shape modeling therefore provides a scalable, flexible tool for both computational biology and material-guided experimental design.

## ACKNOWELDGMENTS

Imaging of the intestinal epithelial cells were performed at the Biofrontiers Institute’s Advanced Light Microscopy Core (RRID: SCR_018302). The Nikon AXR Laser Scanning Confocal is supported by NIH Grant 1S10OD034320.

## FUNDING

We acknowledge funding from the National Institutes of Health F32 HL176073 awarded to A.K., American Heart Association 20PRE35200068 awarded to D.B., and National Institutes of Health R01 HL171197 awarded to KSA.

## AUTHOR CONTRIBUTIONS

A.K. performed image analysis, computational modeling, and generative shape modeling. M.W.Y. and D.B. designed and performed experiments. A.K., M.W.Y., and K.S.A. contributed to writing and editing the paper.

## COMPETING INTERESTS

The authors report no competing interests.

## DATA AND CODE AVAILABILITY

The data utilized in this manuscript is publicly available (https://doi.org/10.25810/degy-jc81). The code used to generate 2D (https://github.com/alexkhang18/Fourier-Series-Shape-Analysis-Fibroblasts) and 3D (https://github.com/alexkhang18/SPHARM-curvature) shape models are available on public repositories.

## Notes

### Competing Interest Statement

The authors have declared no competing interest.

