## Supplementary materials for "Integration and harmonization of cell shape images for generative modeling"

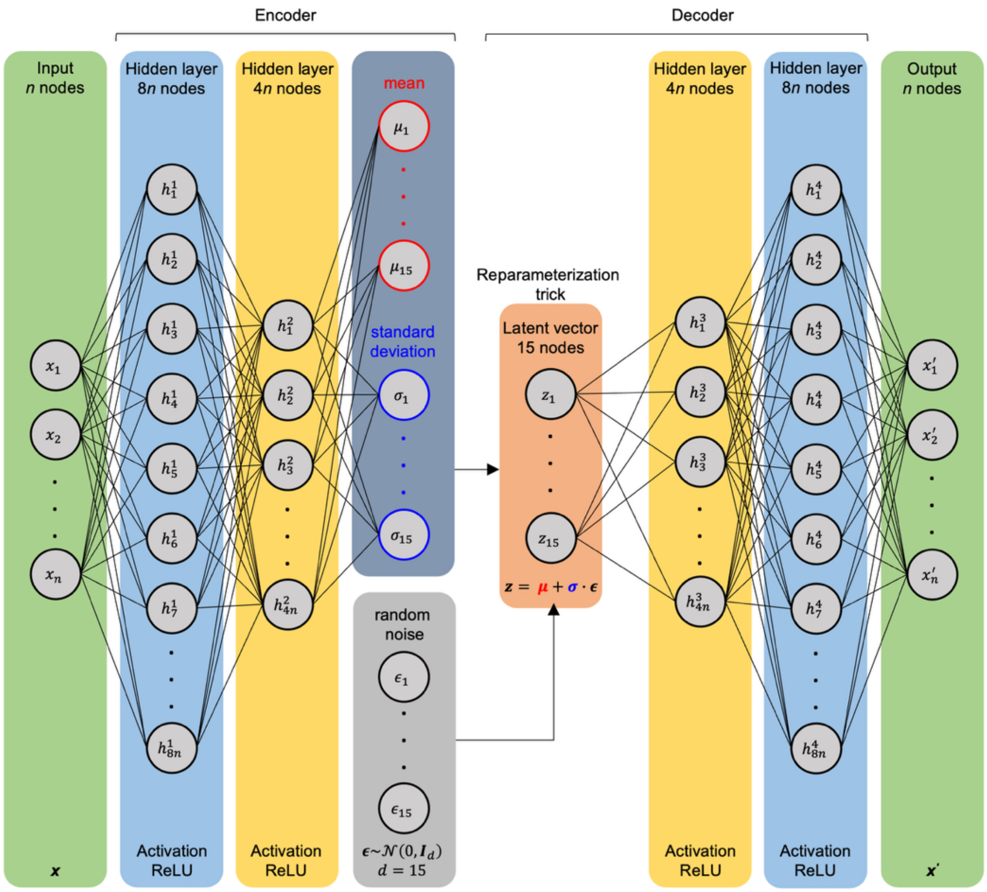


**Fig. S.1 | Architecture of the β-VAE used for generative shape modeling of VICs and IECs.** Separate β-VAEs are trained for each cell type. VIC feature vectors have length n = 87 and are comprised of Fourier series coefficients, DOA, and one-hot encodings for hydrogel stiffness (soft or stiff hydrogel culturing condition). IEC feature vectors have length n = 729 and are comprised of spherical harmonic coefficients, height, and one-hot encodings for cell-type (deformed or non-deformed). The β-VAE is comprised of an encoder with two fully connected and subsequent linear layers with 8*n and 4*n nodes. The encoder outputs mean and standard deviation vectors which are used to compute the latent vector z. The latent vector is then inputted into the decoder whose architecture is the mirror image of the encoder (fully connected subsequent layers with 4*n and 8*n nodes) and results in reconstructed feature vectors.

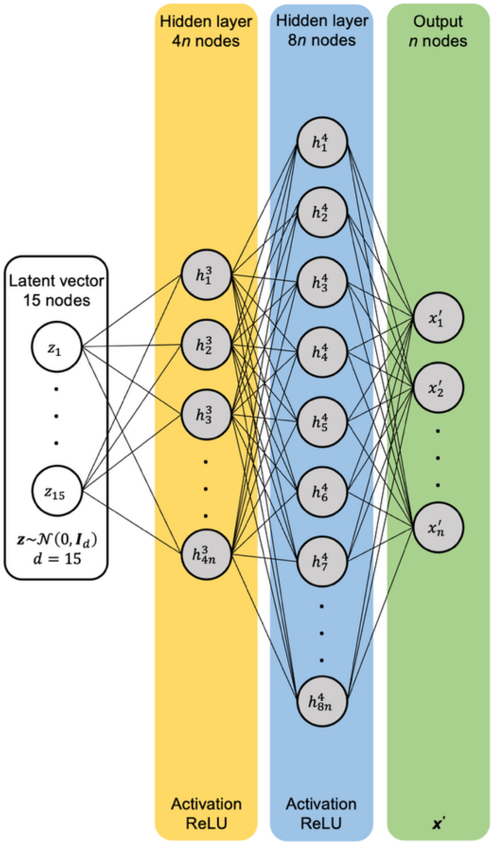


**Fig. S.2 | Generative shape modeling of VICs and IECs are achieved using the trained decoder of their respective β-VAE.** A latent vector of size 15 x 1 is generated by drawing from a standard normal distribution and inputted into the trained decoder to generate unique, but biologically plausible feature vectors. The feature vectors are then used to create cell shape models through either Fourier series or spherical harmonic expansion.

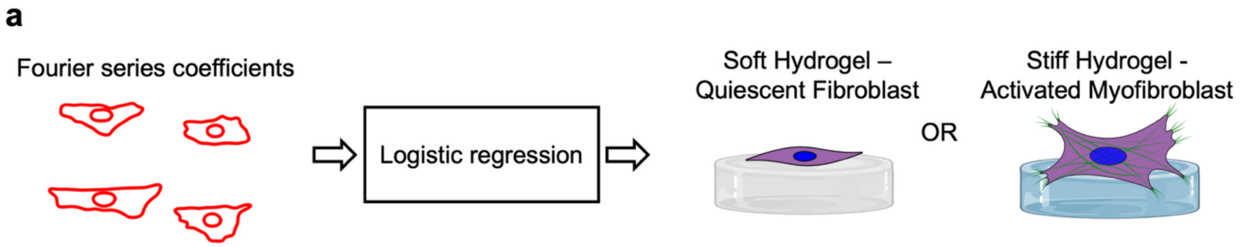


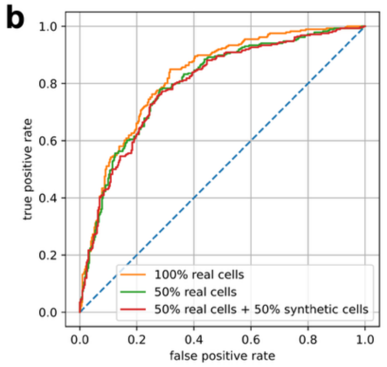


**c**

| Training data | # real cells | # synthetic cells | Accuracy | AUC |
|---|---|---|---|---|
| 100% real data | 2470 | 0 | 75.08% | 0.8237 |
| 50% real data | 1235 | 0 | 73.79% | 0.8006 |
| 50% real data + 10% synthetic data | 1235 | 247 | 73.79% | 0.7991 |
| 50% real data + 20% synthetic data | 1235 | 494 | 74.27% | 0.7993 |
| 50% real data + 30% synthetic data | 1235 | 741 | 74.43% | 0.7965 |
| 50% real data + 40% synthetic data | 1235 | 988 | 73.79% | 0.7965 |
| 50% real data + 50% synthetic data | 1235 | 1235 | 73.79% | 0.7950 |

**Fig. S.3 | Investigating the effect of using synthetically generated data to augment training of logistic regression models for phenotypic grading of VICs. a,** VIC shapes are modeled using Fourier series coefficients which are then used to train a multivariable logistic regression model to classify VICs as being from the soft- or stiff-hydrogel condition. **b,** Receiver operating characteristic curves of models trained on 100% real cells, 50% real cells, and 50% real cells + 50% synthetic cells. The diagonal line denotes the performance of a random classifier. **c,** Summarized accuracy and area under the curve (AUC) of the ROC curve of models trained on real cells or a combination of real and synthetically generated cells. Each model was evaluated using the same held-out test dataset comprised of 618 randomly selected real VICs.

**a**

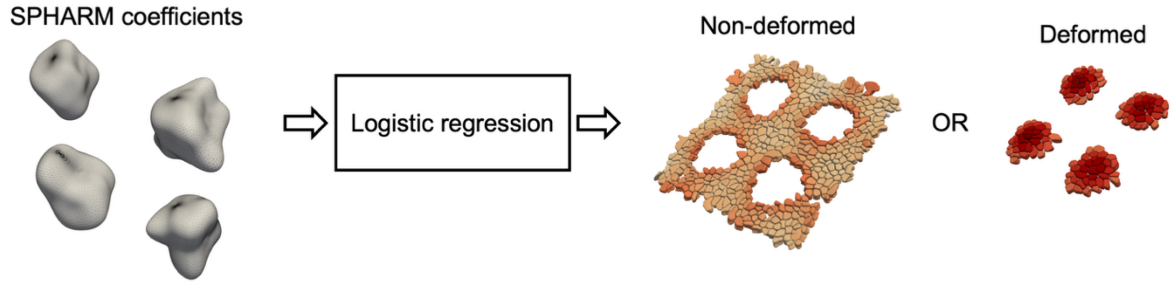


**b**

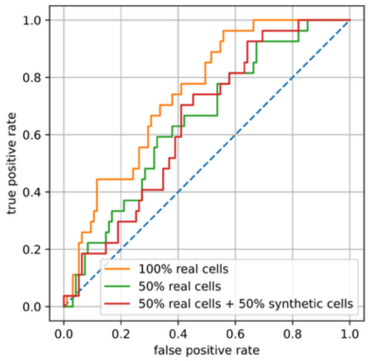


**c**

| Training data | # real cells | # synthetic cells | Accuracy | AUC |
|---|---|---|---|---|
| 100% real data | 486 | 0 | 68.85% | 0.7442 |
| 50% real data | 243 | 0 | 65.57% | 0.6483 |
| 50% real data + 10% synthetic data | 243 | 49 | 65.57% | 0.6581 |
| 50% real data + 20% synthetic data | 243 | 98 | 67.21% | 0.6667 |
| 50% real data + 30% synthetic data | 243 | 146 | 68.85% | 0.6713 |
| 50% real data + 40% synthetic data | 243 | 195 | 66.39% | 0.6581 |
| 50% real data + 50% synthetic data | 243 | 243 | 65.57% | 0.6433 |

**Fig. S.4 | Investigating the effect of using synthetically generated data to augment training of logistic regression models for phenotypic grading of IECs. a,** IEC shapes are modeled using spherical harmonic coefficients which are then used to train a multivariable logistic regression model to classify IECs as being a deformed or non-deformed cell. **b,** Receiver operating characteristic curves of multivariable logistic regression models trained with 100% real cells, 50% real cells, and 50% real cells + 50% synthetic cells. The diagonal line denotes the performance of a random classifier. **c,** Summarized accuracy and area under the curve (AUC) of the ROC curve of models trained on real cells or a combination of real and synthetically generated cells. Each model was evaluated using the same held-out test dataset comprised of 122 randomly selected real IECs.

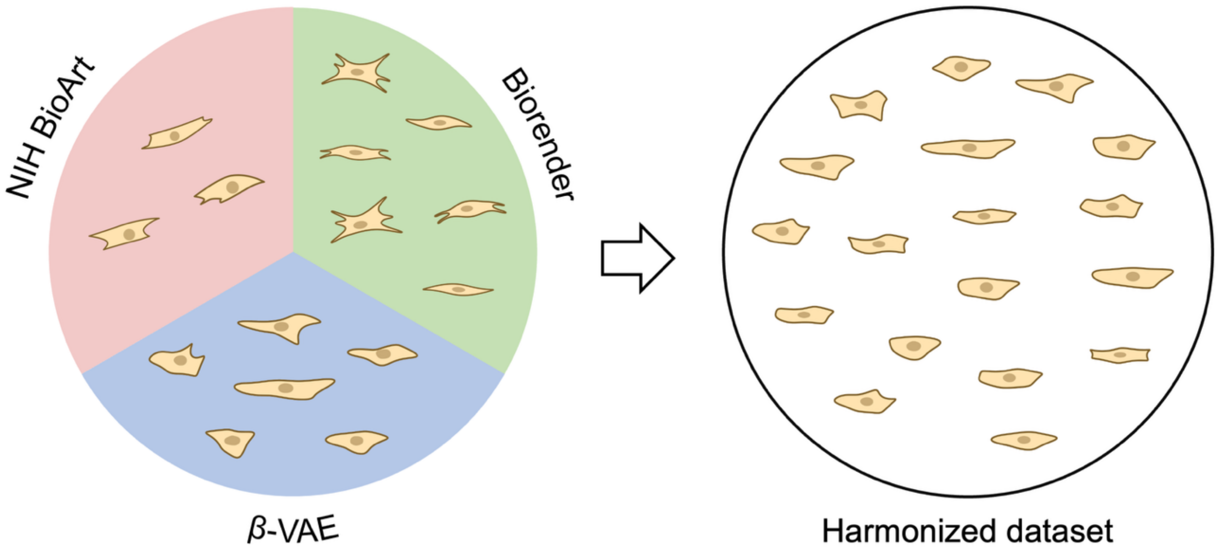


**Fig. S.5 | Harnessing generative modeling of cellular shapes for scientific illustrations.** The VIC-trained β-VAE generate cell illustrations that complement commercial (Biorender) and open-source (NIH BioArt) images. An advantage of using β-VAEs is the ability to generate a limitless number of cells, each with diverse morphological features that are based on real cell shapes. In addition, using cell shape models allow for data harmonization which was accomplished by first subjecting the cell illustrations from NIH BioART and Biorender to Fourier series modeling and then combining them with the VIC dataset to train a β-VAE capable of generating cells that are informed by all three image sources.

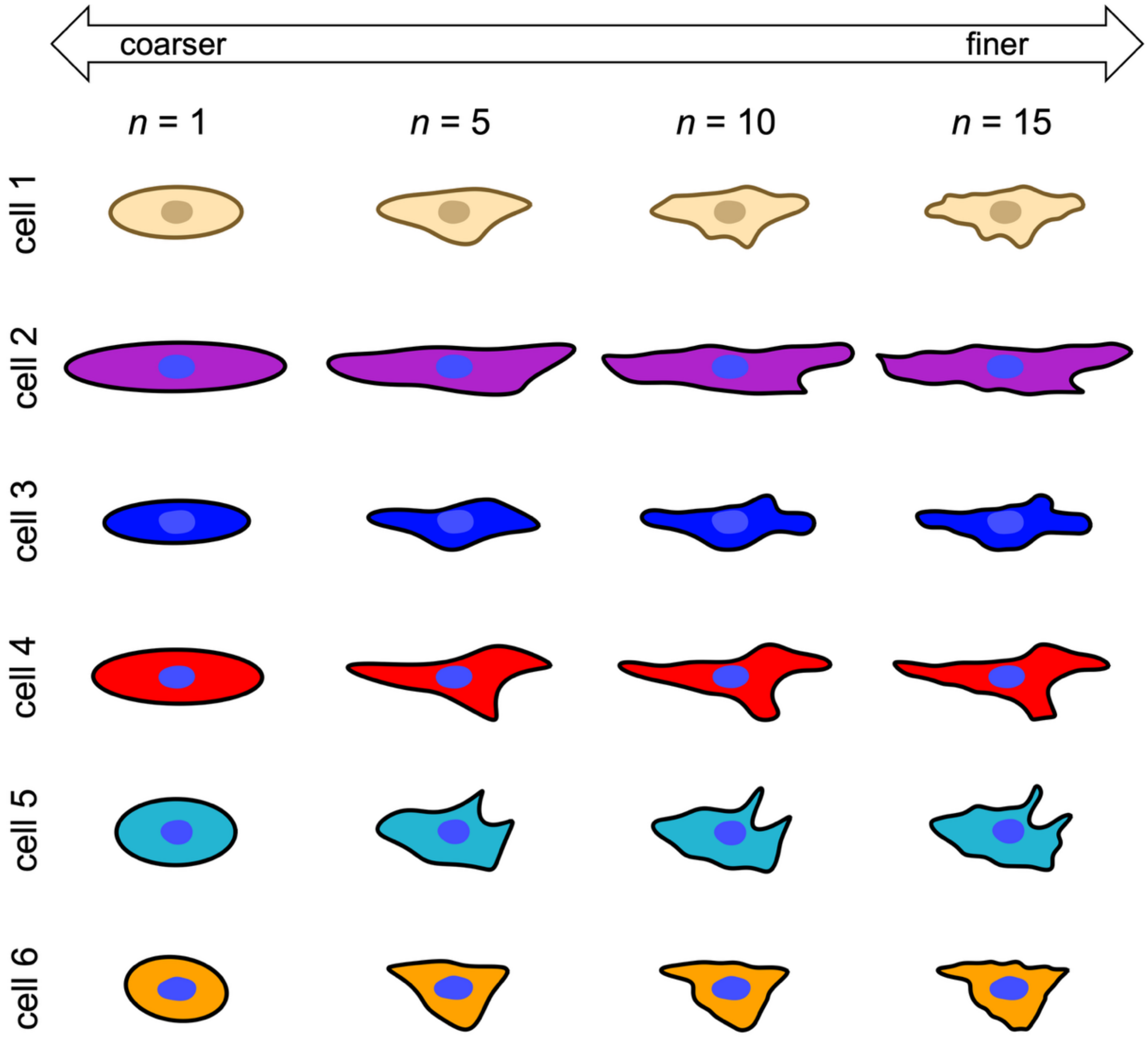


**Fig. S.6 | Generated cell illustrations are highly customizable.** For example, users can choose any color for the stroke and fill of the cell body and nucleus to suit their preferences. In addition, tuning the maximum limit of Fourier series ($n$) gives user control over cell shape coarseness and fineness. Furthermore, using $n$ = 1 gives rise to circular and ellipsoidal cell shapes that can be used to illustrate cells such as blood cells, ova, chondrocytes, and smooth muscle cells.

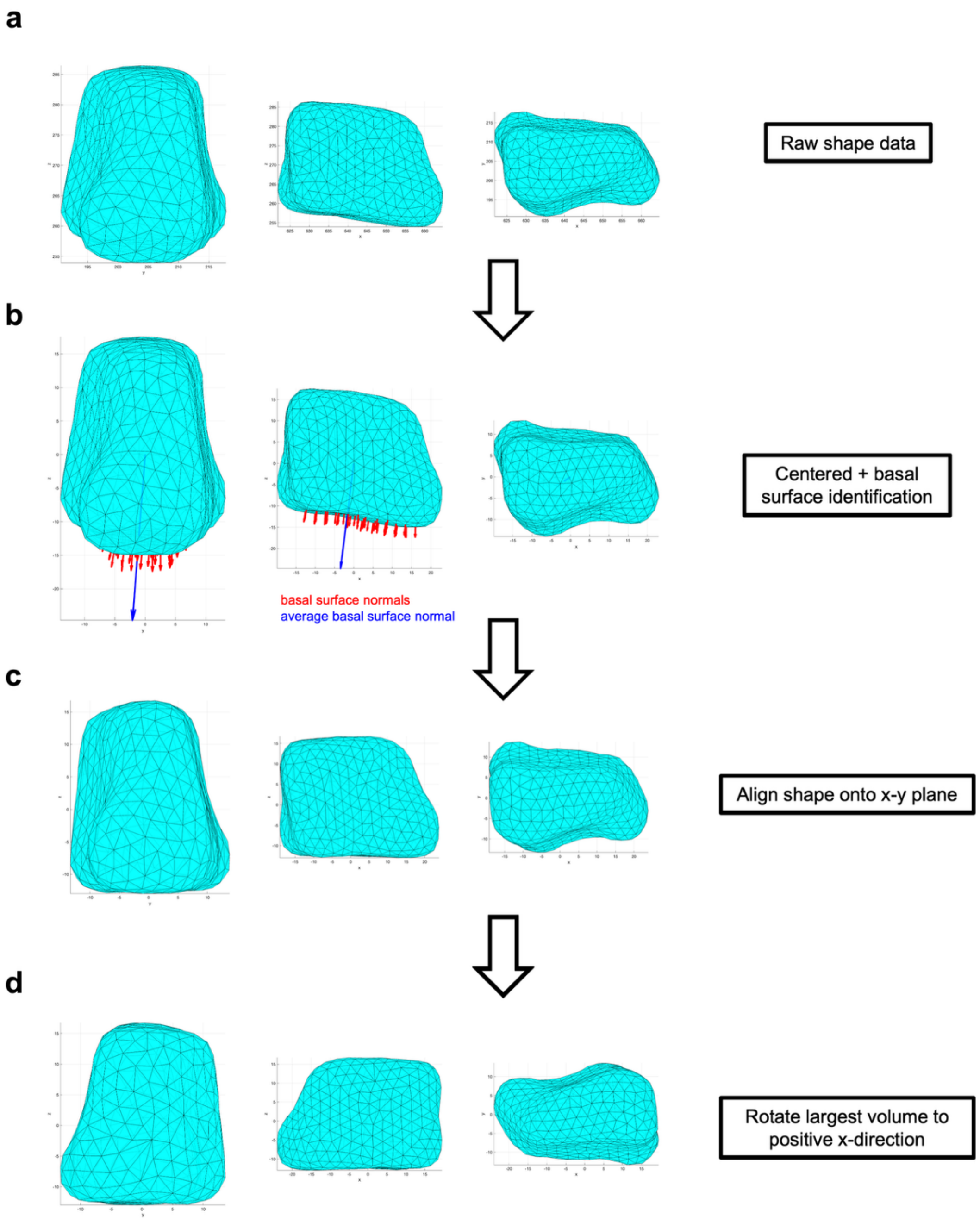


**Fig. S.7 | Registration of IEC surface meshes prior to spherical harmonic shape modeling. a,** Raw 3D IEC surface mesh. **b,** The same mesh in (a) centered on its centroid. The basal surface of the IEC is demarcated by facets with surface normal containing substantial negative values in the global z-direction (red arrows). Averaging the surface normal of the basal surface facets results in an average surface normal (blue arrow). **c,** The 3D surface mesh is rotated so that the average surface normal aligns with the global z-direction (i.e., n = 0, 0, -1), ensuring the basal surface is flat on the x-y plane. **d,** Finally, the shape is rotated in the x-y plane so the largest volume of the cell is located in the positive x direction.

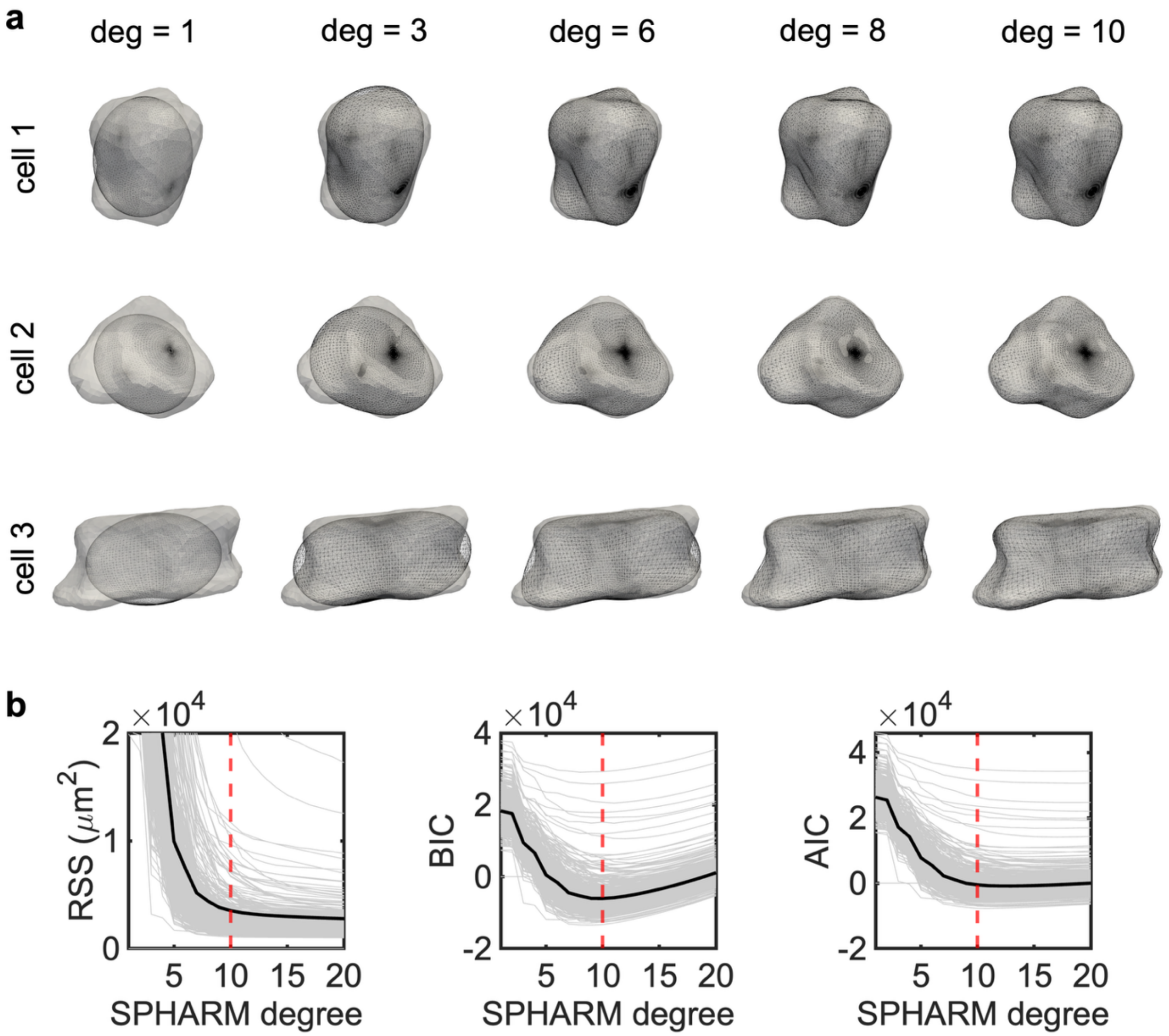


**Fig. S.8 | Spherical harmonic reconstruction of IEC shapes. a,** Three representative IECs modeled using spherical harmonics at varying degrees. **b,** The appropriate number of spherical harmonic degrees was chosen empirically to be 10 for IECs based on the observation in which increasing the number of harmonics no longer produced an appreciable decrease in the residual sum of squares (RSS) and resulted in minimum or near-minimum values for the Bayesian Information Criterion (BIC) and Akaike Information Criterion (AIC). The gray lines denote the results for individual IECs. The black line denotes the mean of the gray lines. The red line denotes the empirical cutoff for the total number of degrees used in spherical harmonic representation of IECs.

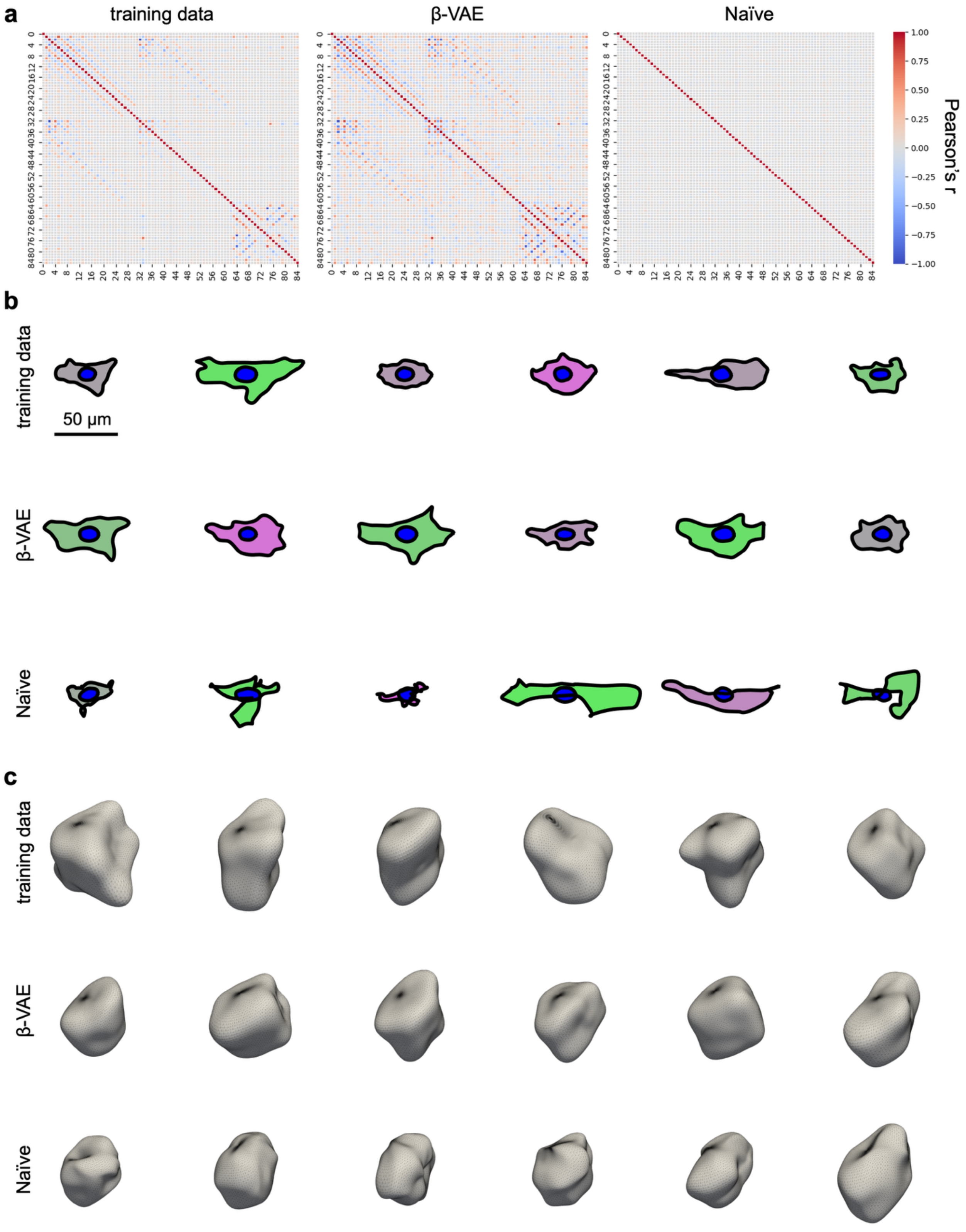


**Fig. S.9 | Comparison of using a β-VAE vs a naïve approach for generative shape modeling.** In the naïve approach, a kernel density estimate is computed for each component of the cell feature vectors.

Random values are then drawn from each kernel density estimate to achieve a synthetic cell feature vector. **a,** Several components of the feature vectors are correlated in the training data, which the β-VAE (but not the naïve approach) can mimic in generated synthetic datasets. Correlation heat maps are provided for the VIC dataset only for brevity. **b,** Representative VIC shape models from the training dataset as well as synthetic VICs generated using the β-VAE and the naïve approach. Note that the β-VAE results in biologically plausible VIC shapes whereas the naïve approach results in nonsensical cell shapes. **c,** Representative IEC shape models from the training dataset as well as synthetic IECs generated using the β-VAE and the naïve approach. Note that the β-VAE results in biologically plausible IEC shapes whereas the naïve approach results in non-smooth IECs with unrealistic grooves and dimples.

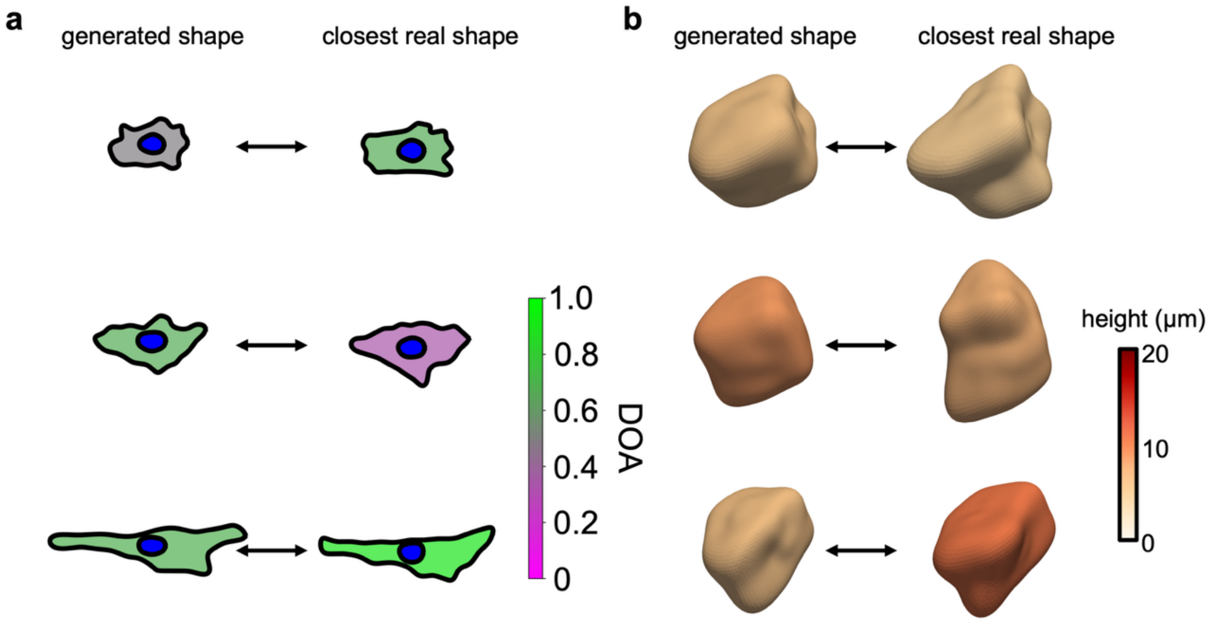


**Fig. S.10 | β-VAEs generate unique but biologically plausible cells and do not simply reproduce the training dataset. a,** Three representative VICs generated using a β-VAE and their closest match to real cells in the training dataset. **b,** Three representative IECs generated using a β-VAE and their closest match to real cells in the training dataset.
